## Supplementary material for "Power-law adaptation in the presynaptic vesicle cycle": SI

**This file includes**

Supplementary figures S1-S18

Supplementary table 1

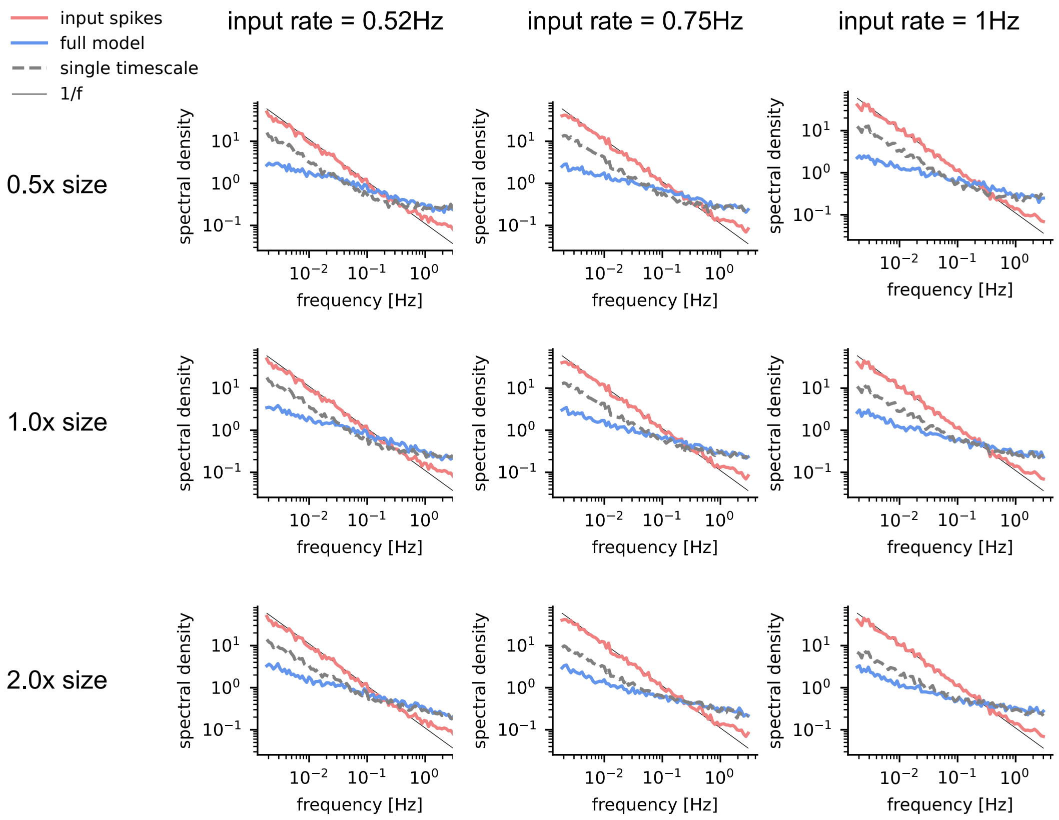


**Fig S1. Whitening is robust to changes in synaptic size and rate of input.** We performed the same experiment as in the main paper (Fig 4B), but varied the average rate of the input spike-train, and the size of the synapse (by scaling the maximum number of vesicles per pool, but leaving the timescales as before). Partial whitening stays intact also when changing the sizes of all vesicle pools by 0.5 or 2 fold, and when changing the total rate of 1/f input spike trains.

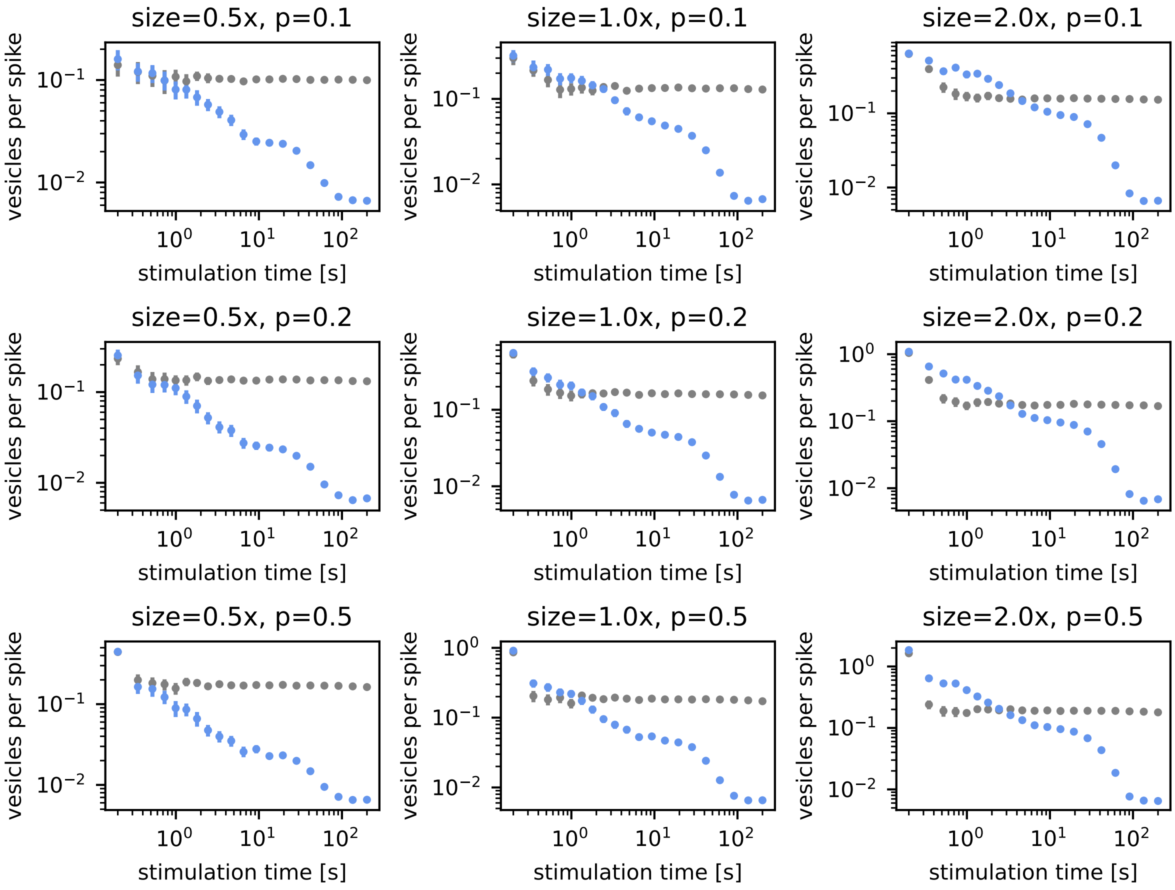


**Fig S2. Detailed dynamics of depression depend on synapse size and release probability (Stimulation rate = 50Hz).** Change of released vesicles per spike under constant stimulation of 50Hz as in the main paper (Fig 2B). The detailed depression dynamics depend on synapse size (scaling the maximum number of vesicles per pool) and release probability. Still, all full models (blue) show ongoing depression over several orders of magnitude of stimulation time. The single timescale model (gray) does not show this behavior.

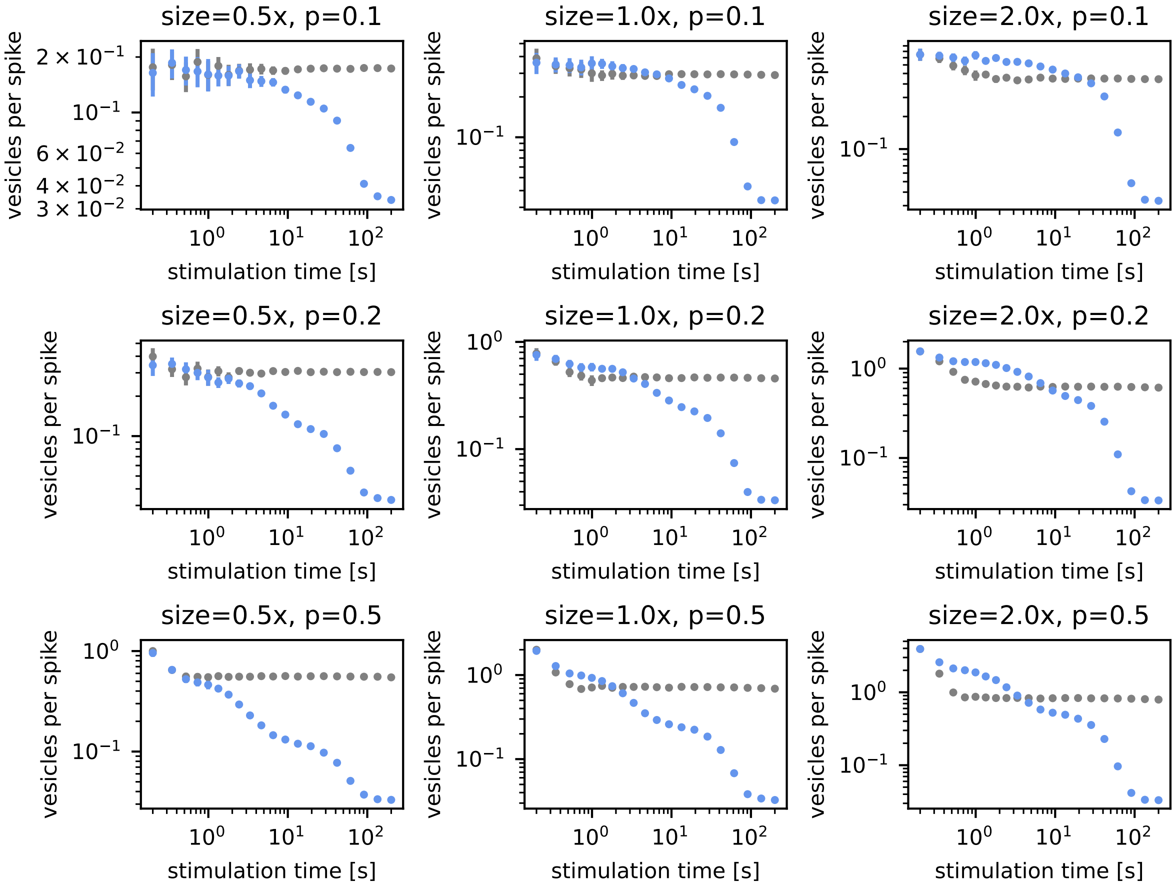


**Fig S3. Detailed dynamics of depression depend on synapse size and release probability (Stimulation rate = 10 Hz).** Change of released vesicles per spike under constant stimulation as in the main paper (Fig 2B) but with slower stimulation rate of 10Hz. Again, the detailed depression dynamics depend on synapse size and release probability. All full models show ongoing depression over several orders of magnitude of stimulation time. Note, that when all pools are exhausted the recovery rate becomes the bottleneck, after which the released vesicles per spike decrease to a baseline. This results in a less clearly visible power-law depression for large synapses with low release probabilities.

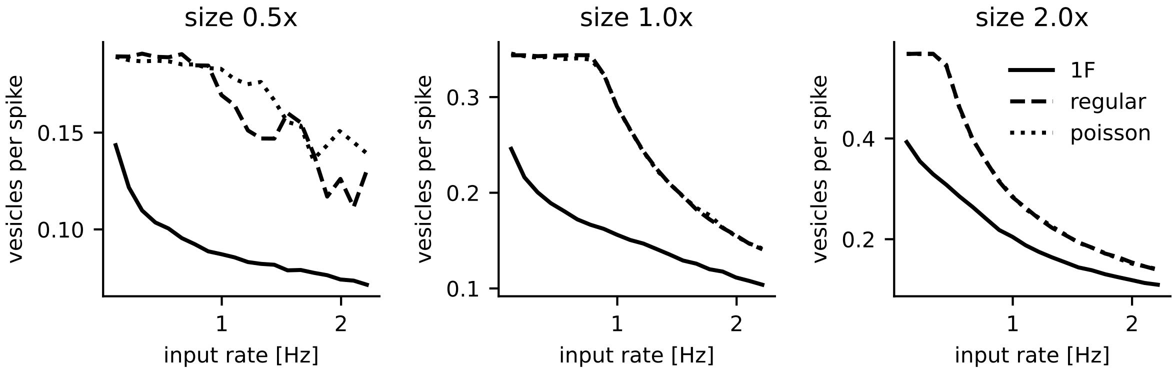

**Fig S4. The synapse model reacts nonlinearly to stimulation.** Depending on the stimulation protocol (1/f spiking, spikes with regular intervals, Poisson spiking) the input-output rate function of the synapse is different. When the synapse starts depressing, the released vesicles per spike start to decrease. For 1/f spiking this happens earlier, since here rapid bursts of presynaptic spikes can occur also for low average input rate.

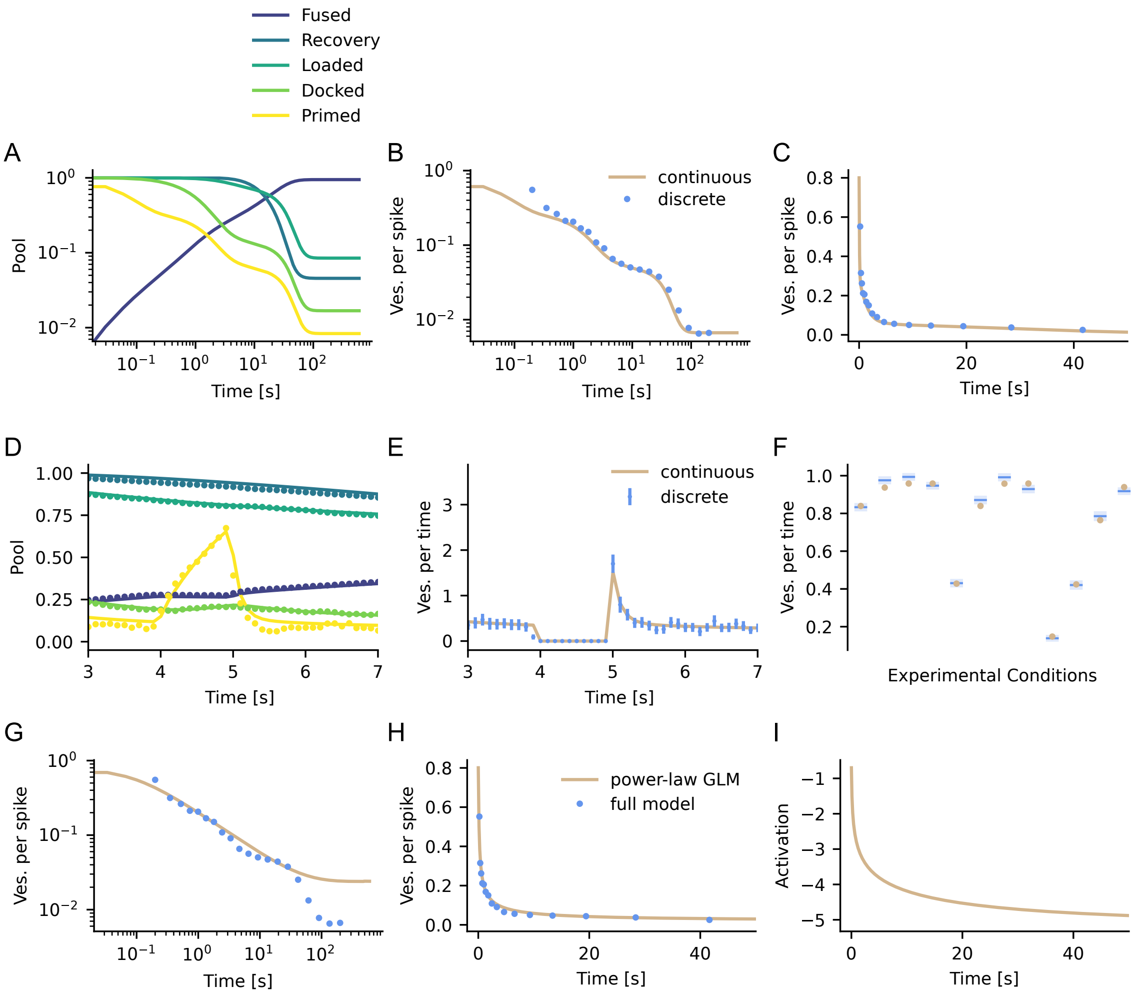


**Fig S5. Approximate continuous solution to vesicle release dynamics. A,B,C** Comparison of depression under constant 50 Hz stimulation averaged over 100 repetitions. **A** Pool dynamics. **B,C** Comparison of vesicle release in the discrete stochastic and the continuous model under 50 Hz stimulation, in log-log (B) and regular (C) scale. Note, that the first bin of the discrete model takes into account all releases before that and hence is biased upwards compared to the continuous model. **D,E,F** More detailed comparison under stimulation as used in the experiment (Fig 3 in the main paper). Here, discrete dynamics were averaged over 100 realizations in bins of 100 ms. **D** Comparison of discrete and continuous pool dynamics. **E** Comparison of discrete and continuous vesicle release rates. Errorbars denote 95% confidence intervals computed from error of the mean. **F** Comparison of discrete and continuous results for the experimental paradigm (Fig 3 in the main paper), i.e., average over 2 seconds 20 Hz stimulation after exhaustion and pause period. Conditions are the same as in the main paper figure 3. Errorbars denote 95% confidence intervals computed from error of the mean. Shaded areas denote standard deviation from mean. **G,H** Releases per spike comparison (as in B,C) of the discrete model with the fitted GLM (Fig S16), where the parameters are $b = -0.7$, $\alpha = 0.3$ and $\kappa_0 = 0.1$. Note, that the linear model generalizes to the new data under normal depression, but does not fit the nonlinear behavior of the synapse when it is strongly depressed to the point of failure. For plotting we here used the implementation of the model as described in Fig S14. **I** Associated activation $a(t)$ (Eq 3) over time.

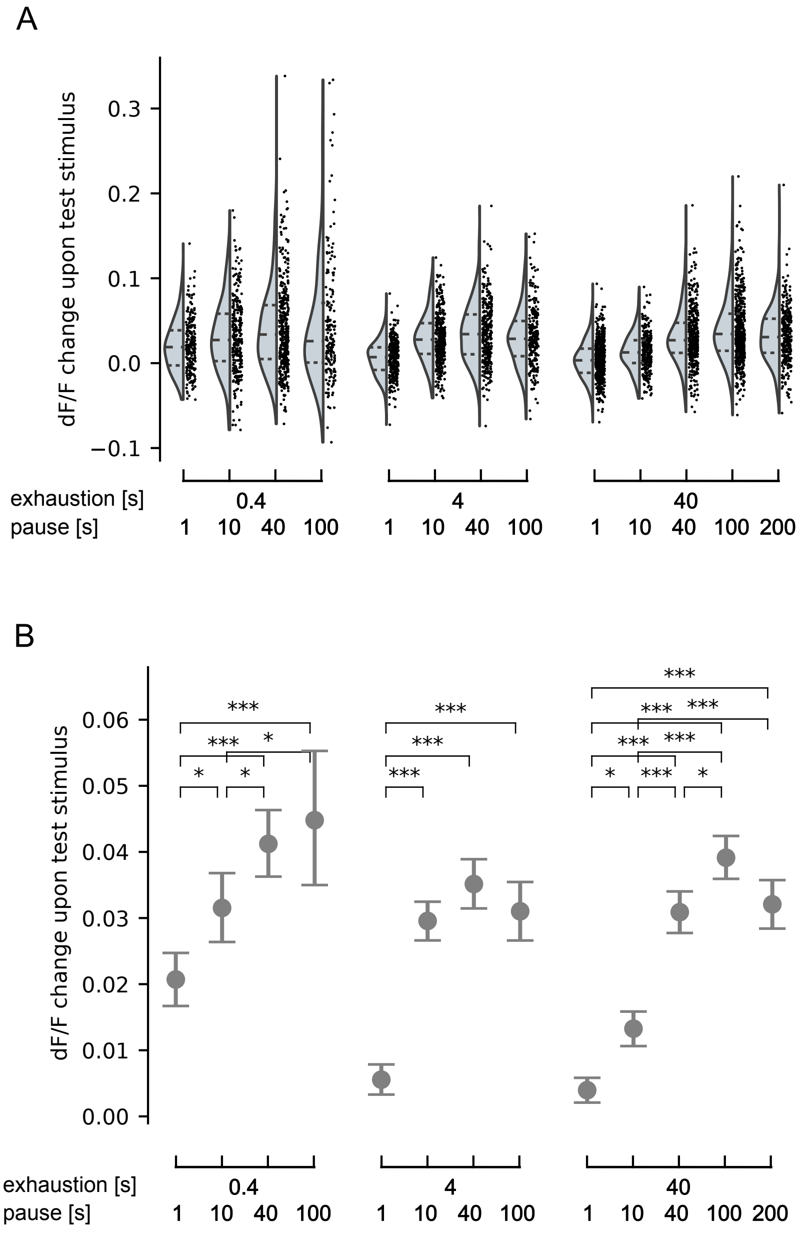


**Fig S6. Summary of experimental data. A** Violin plot showing medians and quartiles, and the associated data points. The number of data-points in each condition are (from left to right) 229, 283, 409, 186, 321, 375, 343, 244, 547, 279, 419, 519, 308. **B** Mean data values with 95% bootstrap confidence intervals, as in the main manuscript. Conditions were compared using the Kruskal-Wallis test, followed by the Tukey post-hoc test, where * $p < 0.05$, ** $p < 0.005$, *** $p < 0.0005$.

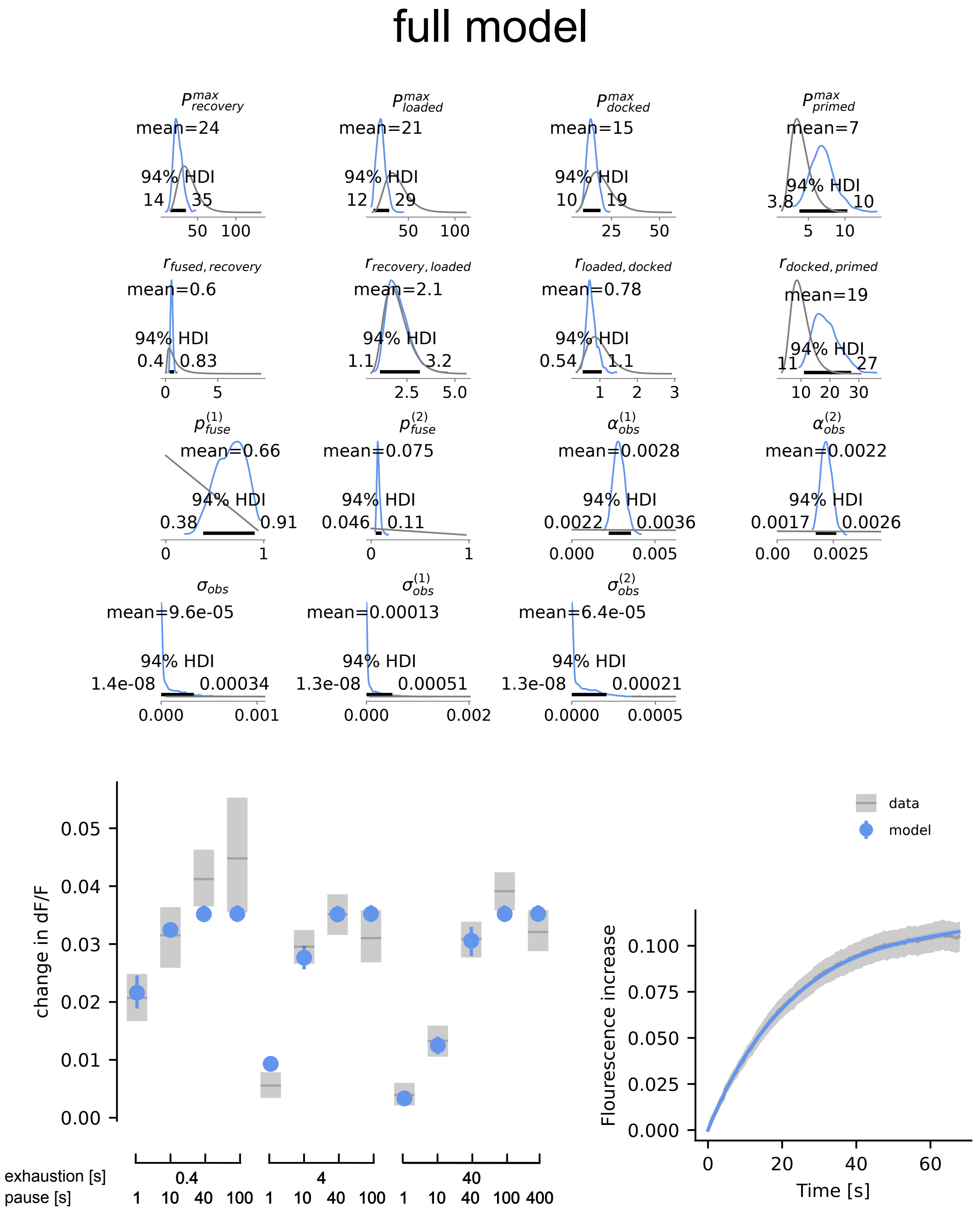


**Fig S7. Summary of the full model MCMC posterior.** Top: Prior (gray) and posterior (blue) distributions of model parameters. Bottom: Comparison of data and model observations posterior, as in the main text.

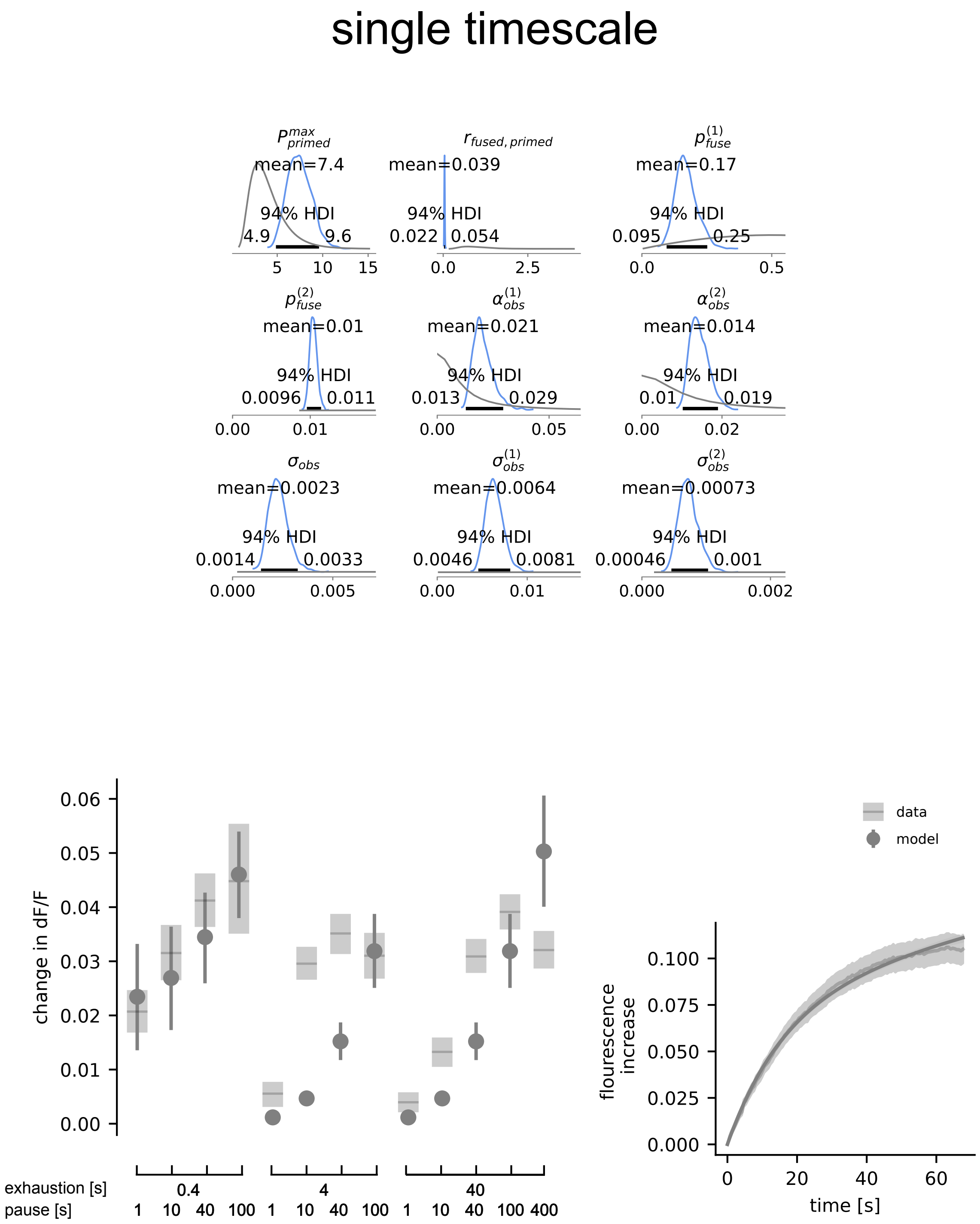


**Fig S8. Summary of the single timescale model MCMC posterior.** Top: Prior (gray) and posterior (blue) distributions of model parameters. Note, that here for $p_{fuse}$ we chose a Beta(2,2) prior instead of a Beta(2,1) prior, to prevent extremely small posterior estimates. Still, release probabilities that allow to model the experimental results are very small. Bottom: Comparison of data and model observations posterior, as in the main text.

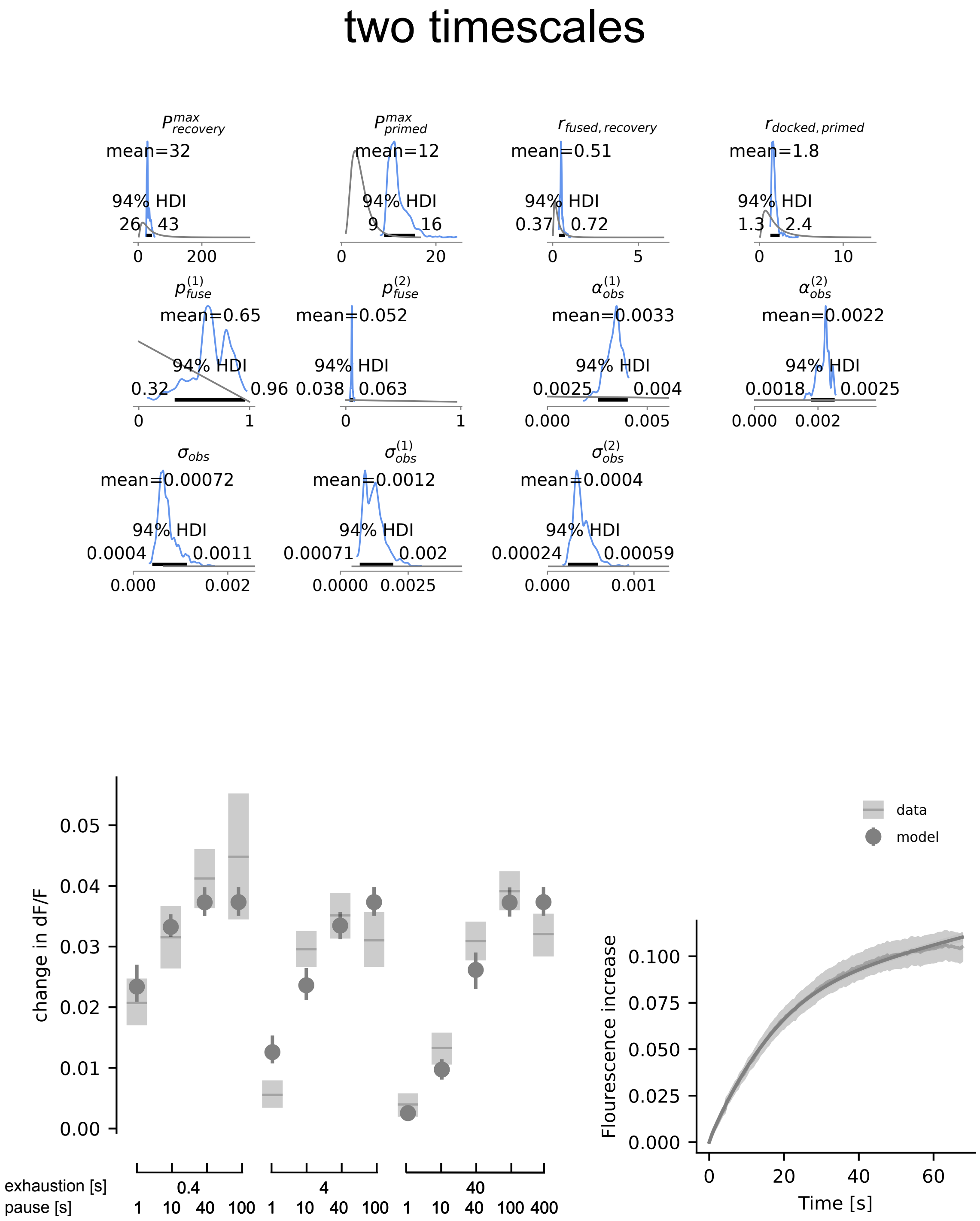


**Fig S9. Summary of the two timescale model MCMC posterior.** Top: Prior (gray) and posterior (blue) distributions of model parameters . Bottom: Comparison of data and model observations posterior, as in the main text.

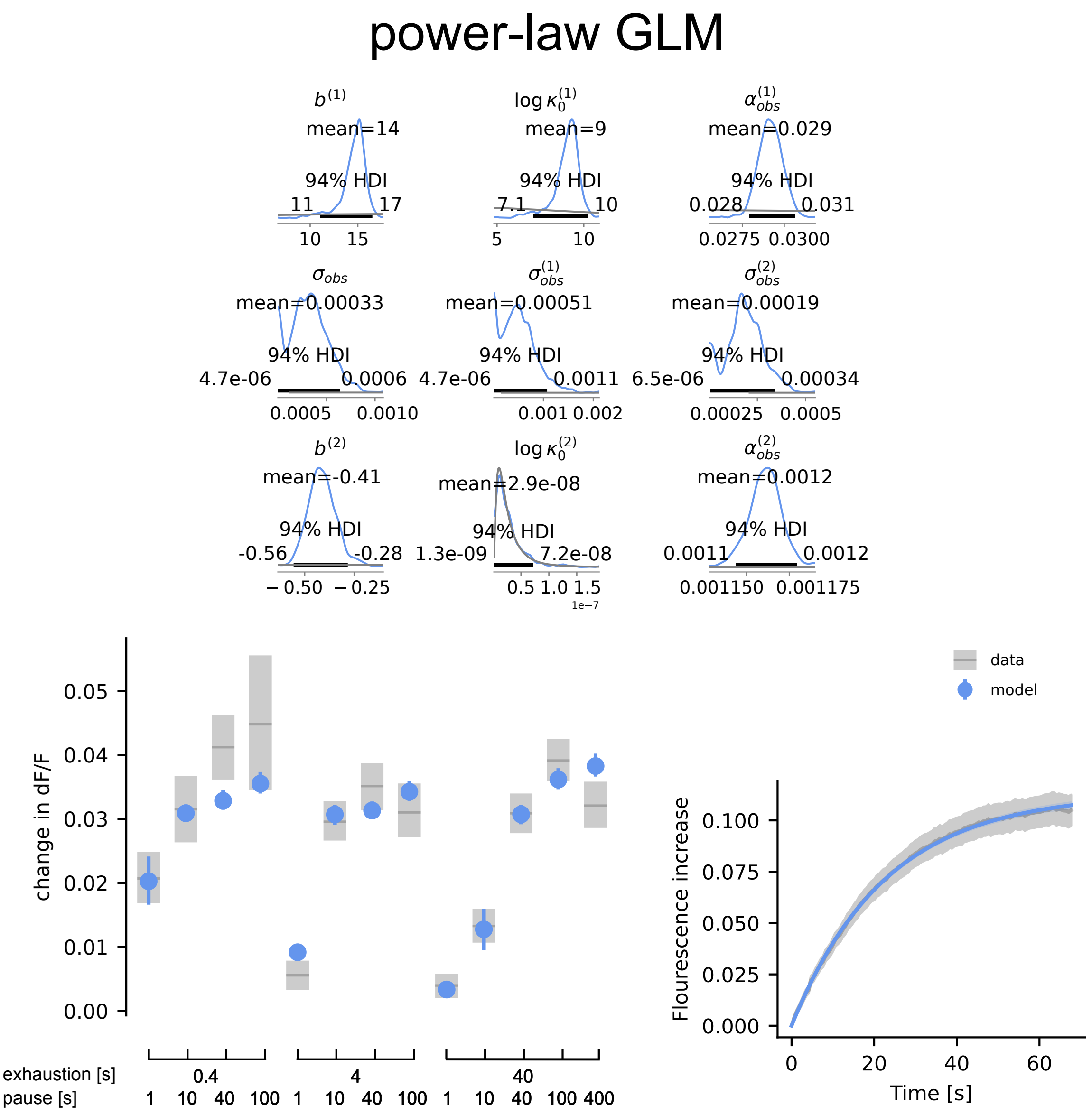


**Fig S10. Summary of the MCMC posterior for a model based on the GLM.** This model is a continuous-release variant of the GLM, where the sum over spikes in equation 3 becomes a sum over time. The output of the model is then computed as the expected number of released vesicles $r(t) = \mathbb{E}[k(t)] = \sum_{i=1}^{N} kp(k(t)|\vec{t}_{\text{past}}, \theta)$, and the kernel was parametrized as a truncated power-law $\kappa(\Delta t) = \kappa_0 t^{-\alpha} \exp(-t/T)$, where the cutoff $T = 120$ s. After the cutoff we also set $\kappa(\Delta t) = 0$, for efficiency. The exponent was set to $\alpha = 0.3$, as in the model (Fig S17). To account for changes in Calcium levels in the experiments, two different baseline release rates $b$ and kernel scalings $\kappa_0$ were used. This is necessary, since in this model the single vesicle release probability is not a single parameter, but entangled with the number of resources and stochasticity of the synapse. Top: Prior (gray) and posterior (blue) distributions of model parameters (except $\alpha_{\text{obs}}$, which only shows the posterior since it uses an improper (flat) prior). Bottom: Comparison of data and model observations posterior, as in the main text.

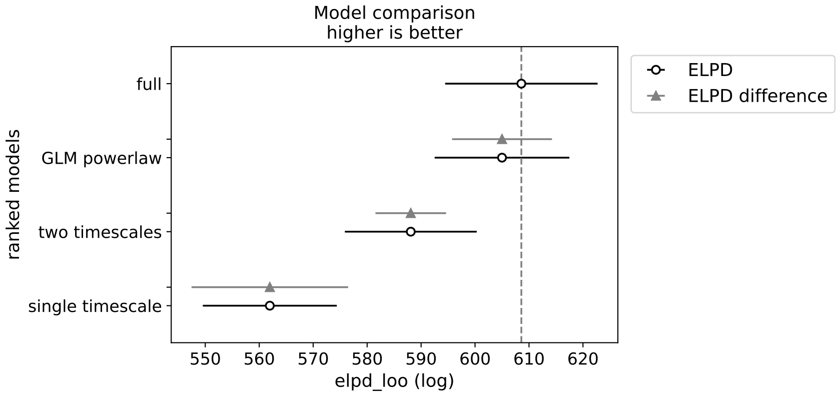


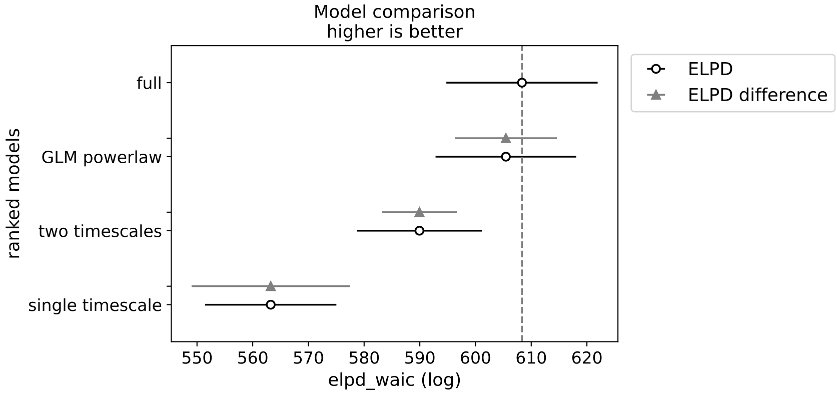


**Fig S11. Model comparison of full, single timescale, two timescales, and power-law model** Model comparison computed using leave-one-out (loo) cross validation (top) and the widely applicable information criterion (WAIC, bottom) as implemented by PyMC3 (see Vehtari, A., Gelman, A., & Gabry, J. (2017). Practical Bayesian model evaluation using leave-one-out cross-validation and WAIC. *Statistics and computing.*). The comparison is based on the expected log pointwise predictive densities (ELPDs), i.e., the expected accuracy on a new dataset, as estimated via WAIC or loo. Plotted are the mean and standard error of the ELPD, and the standard error of the ELPD difference to the best model. In both WAIC and loo, the single timescale, the two timescale, and the power-law model are clearly worse at fitting the data than the multi-timescale model, which is indicated by the standard error of difference in ELPD. It is furthermore possible to assign 'weights' to the models, which can be thought of as estimated probabilities of a model being the best model on new data. Assigned model weights via both loo and WAIC were 1 for the full model and 0 for the other models, again favoring the full model. For more details on the interpretation of these measures, see McElreath, R. (2016). Statistical Rethinking: A Bayesian Course with Examples in R and Stan. *CRC Press.*

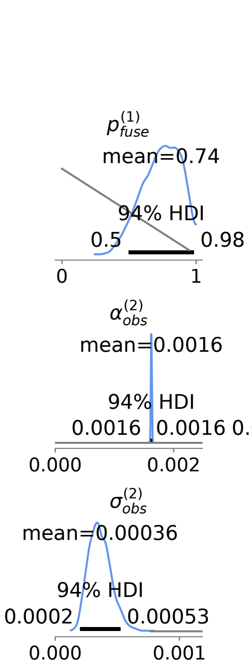


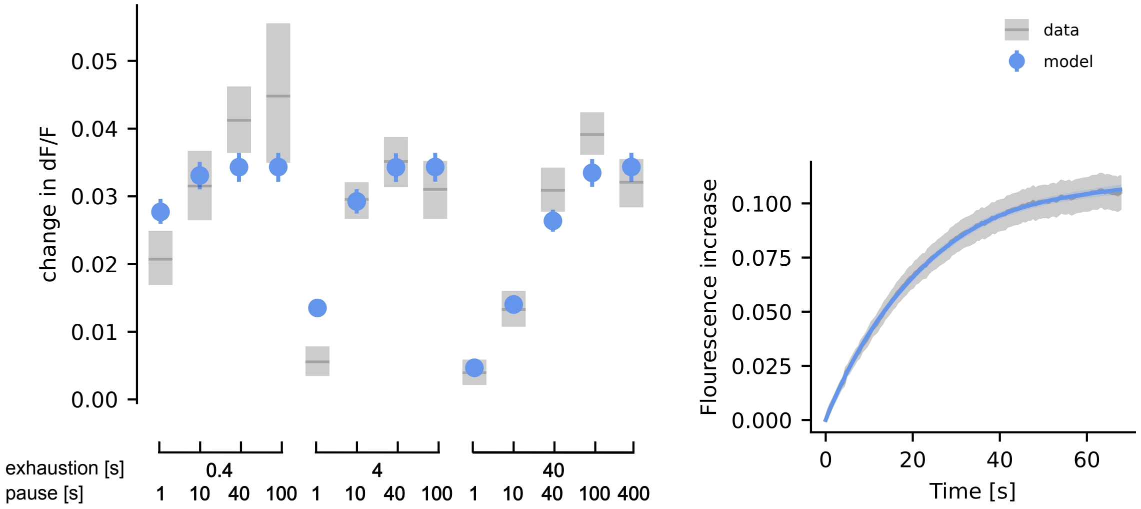


**Fig S12. Summary of the MCMC posterior for a full model where only $p_{fuse}$ is fitted.** Top: Prior (gray) and posterior (blue) distributions of model parameters. Bottom: Comparison of data and model observations posterior, as in the main text.

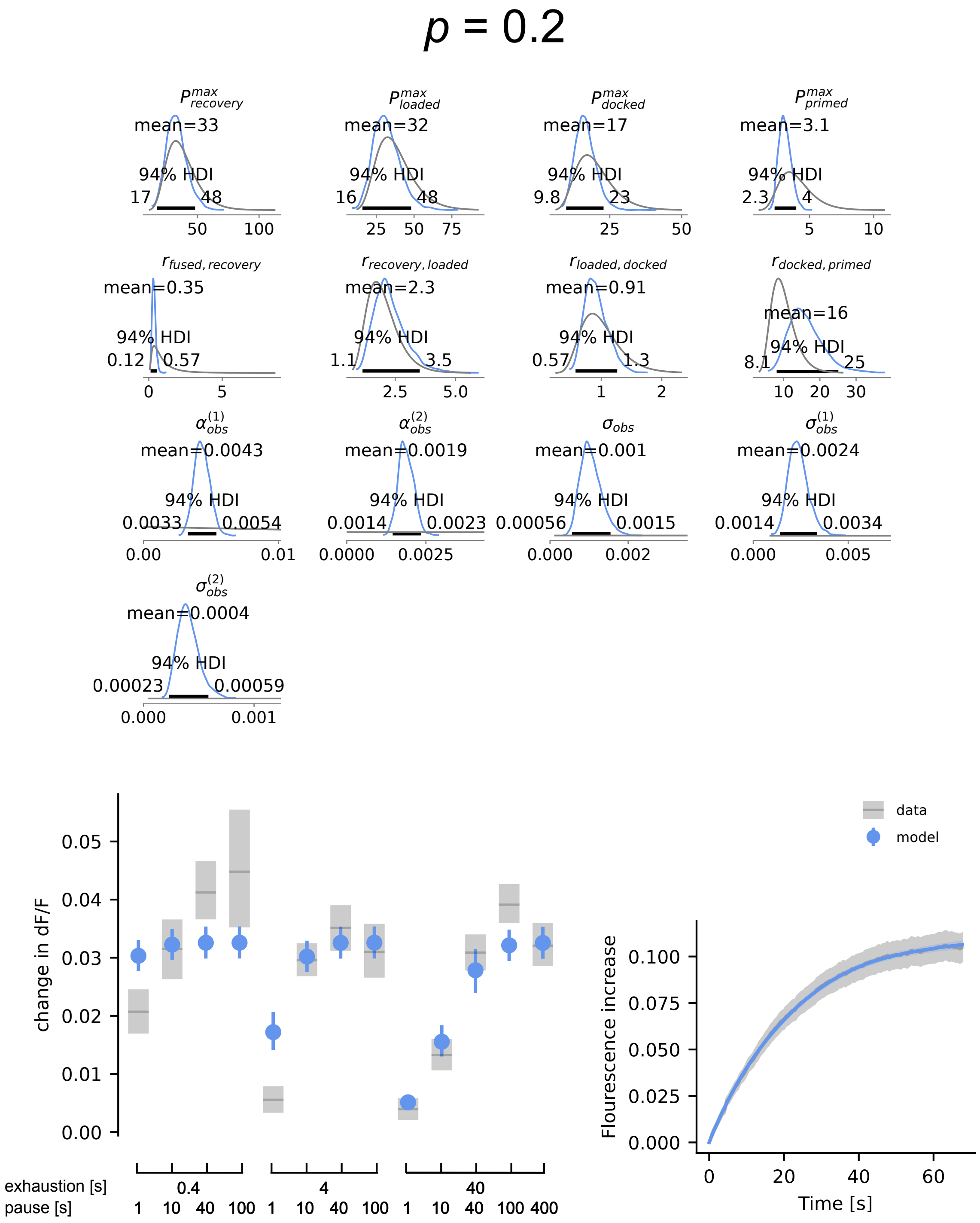


**Fig S13. Summary of the MCMC posterior for a full model where $p_{fuse}$ is set to 0.2.** Top: Prior (gray) and posterior (blue) distributions of model parameters. Bottom: Comparison of data and model observations posterior, as in the main text.

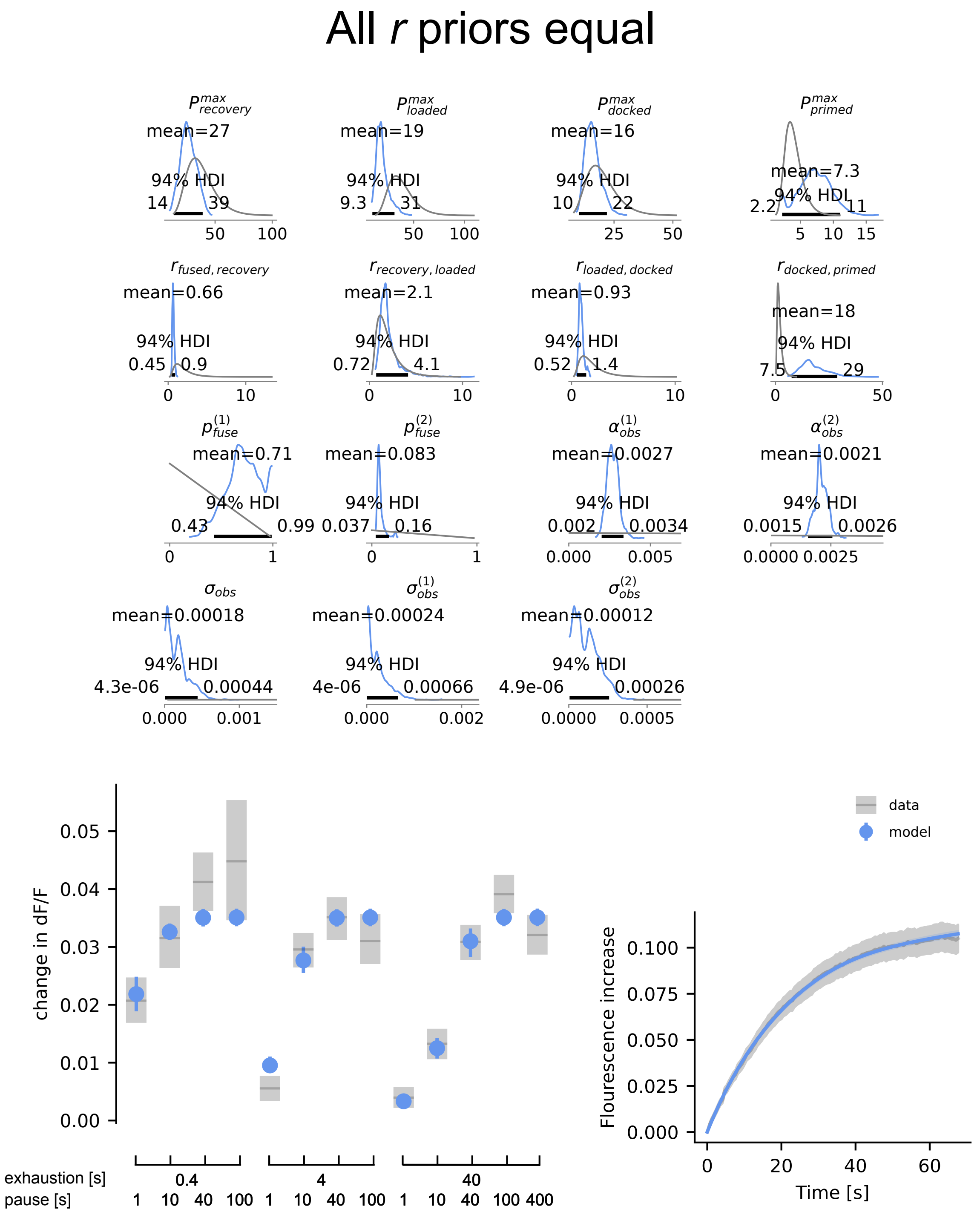


**Fig S14. Summary of the MCMC posterior for a full model where all priors on rates $r$ are set to the same broad distribution.** Top: Prior (gray) and posterior (blue) distributions of model parameters. Bottom: Comparison of data and model observations posterior, as in the main text. Intriguingly, the data constrains the rates well and recovers the same posterior distributions as the well constrained model (Fig S7), which are close to the experimentally informed prior distributions.

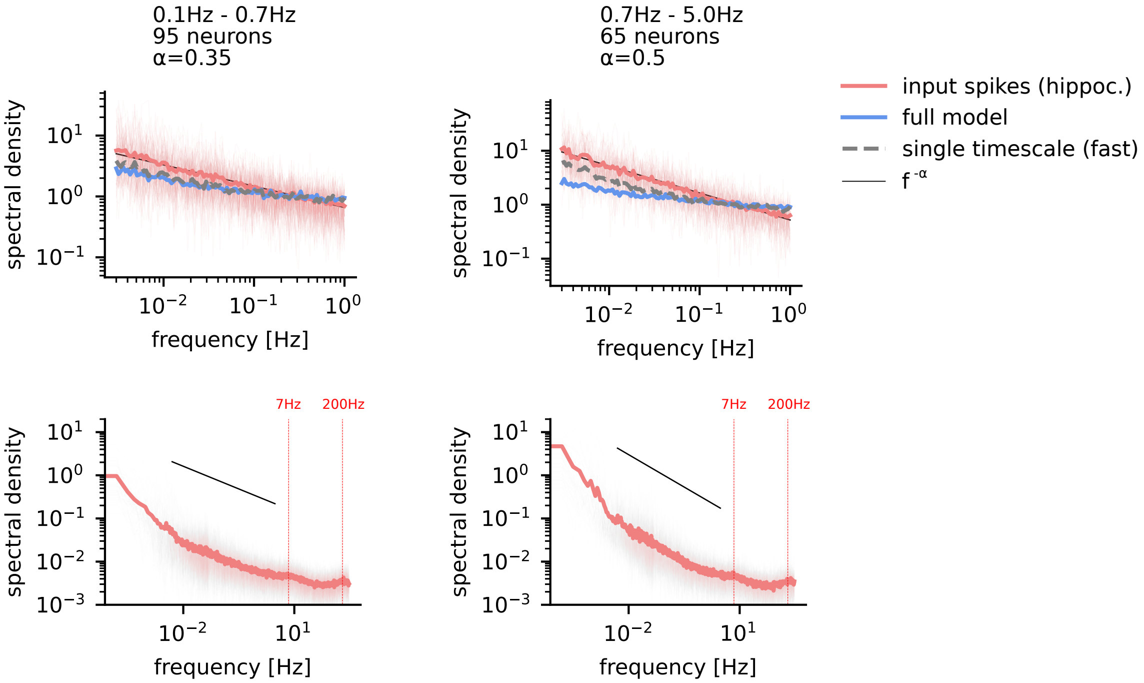


**Fig S15. Whitening results for different hippocampal neuron rates.** The spectral densities of hippocampal neurons show a frequency dependence, with lower rate neurons having a flatter spectrum. (Top) Power law adaptation in the vesicle cycle results in approximate whitening, i.e., a flattening of the input spectrum, for both, high and low rates. However, the difference between the single timescale and the full model is less pronounced for low rate input, where efficient coding might be less important. (Bottom) The power law is present over $\sim 3$ orders of magnitude, as demonstrated above. Small peaks can be observed at 7 Hz (Theta) and 200 Hz.

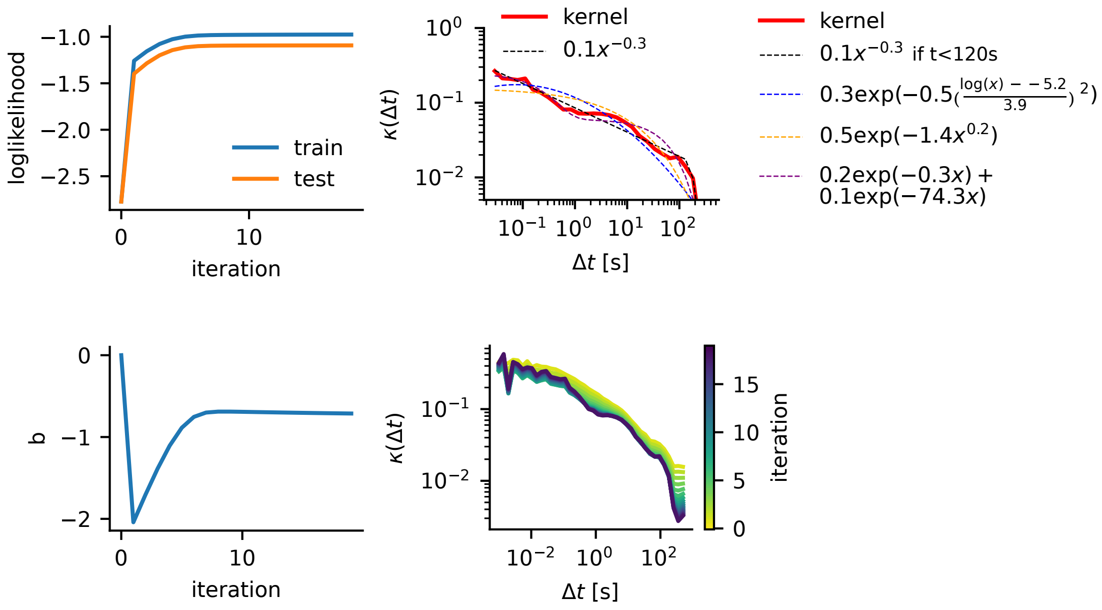


**Fig S16. Validation of GLM fitting.** The maximum likelihood fitting of parameters $b$ and $\kappa(\Delta t)$ converges after about 20 iterations. The likelihood computed on the training and a test set show similar behavior during learning, indicating that the kernel is not overfitting the data. We furthermore tested whether the kernel can be described better by an alternative simple functional form, and fitted also log-normal, exponential (not shown) and stretched exponential functions. The power-law description is favored by both AIC (-101.6, -75.0, -31.8, -33.0, -45.9) and BIC (-97.8, -71.2, -28.6, -30.7, -42.8) for the power-law, double exponential, log-normal, exponential, and stretched exponential, respectively.

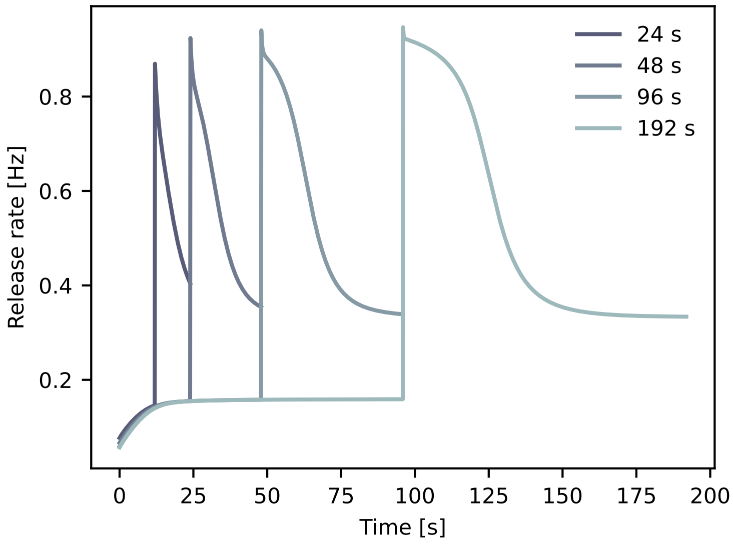


**Fig S17. Demonstration of dynamically adjusted depression rate.** The continuous vesicle cycle model (Fig S5) is stimulated using periodic high-low inputs (high = 1.2 Hz, low = 0.2 Hz). Depending on the period the depression is slower or faster. Compare "Efficiency and ambiguity in an adaptive neural code" by Fairhall et al. in *Nature* (2001).

| Parameter | Value | Distribution |
|---|---|---|
| $r_{\text{fused,recovery}}$ | $0.33\,\text{s}^{-1}$ | LogNormal($\mu = 0.33\,\text{s}^{-1}, \sigma = \mu/3$) |
| $r_{\text{recovery,loaded}}$ | $2\,\text{s}^{-1}$ | LogNormal($\mu = 2\,\text{s}^{-1}, \sigma = \mu/3$) |
| $r_{\text{loaded,docked}}$ | $1\,\text{s}^{-1}$ | LogNormal($\mu = 1\,\text{s}^{-1}, \sigma = \mu/3$) |
| $r_{\text{docked,primed}}$ | $10\,\text{s}^{-1}$ | LogNormal($\mu = 10\,\text{s}^{-1}, \sigma = \mu/3$) |
| $P^{max}_{\text{recovery}}$ | 40 | LogNormal($\mu = 40, \sigma = \mu/3$) |
| $P^{max}_{\text{loaded}}$ | 40 | LogNormal($\mu = 40, \sigma = \mu/3$) |
| $P^{max}_{\text{docked}}$ | 20 | LogNormal($\mu = 20, \sigma = \mu/3$) |
| $P^{max}_{\text{primed}}$ | 4 | LogNormal($\mu = 4, \sigma = \mu/3$) |
| $p_{\text{fuse}}$ | 0.2 | Beta($\alpha = 1, \beta = 2$) |
| $\sigma_{obs}$ | - | HalfCauchy($\beta = 0.2$) |
| $\sigma^{(1)}_{obs}$, $\sigma^{(2)}_{obs}$ | - | Normal($\mu = \sigma_{obs}, \sigma = \sigma_{obs}/4$) |
| $\alpha^{(1)}_{obs}$ ,$\alpha^{(2)}_{obs}$ | - | HalfCauchy($\beta = 0.01$) |

**Table 1.** Summary of parameters values used in the full model (Values) and priors on the Bayesian model parameters (Distribution). Values were taken from Jähne et al, Cell Reports, 2021. Further details about the other models and priors can be found in the simulation code.
